# Compromised CDK12 activity causes dependency on the non-essential spliceosome components

**DOI:** 10.1101/2021.12.07.470703

**Authors:** Satu Pallasaho, Aishwarya Gondane, Damien Duveau, Craig Thomas, Massimo Loda, Harri M Itkonen

## Abstract

Prostate cancer (PC) is the most common cancer in men and after development of the castration-resistant PC (CRPC), there are no curative treatment options. Inactivating mutations in cyclin-dependent kinase 12 (CDK12) define an aggressive sub-type of CRPC. We hypothesized that compromised CDK12 activity leads to significant rewiring of the CRPC cells, and that this rewiring results in actionable synthetic lethal interactions.

**Methods:** We used combinatorial lethal screening, ChIP-seq data, RNA-seq data, global alternative splicing analysis, and comprehensive mass spectrometry (MS) profiling to understand how the compromised CDK12 activity rewires the CRPC cells. In addition, we used DepMap-, PC- and CRPC-datasets as a strategy to identify factors that are selectively required by the CDK12-mutant cells.

**Results:** We show that inhibition of O-GlcNAc transferase (OGT) and CDK12 induces cancer cell-selective growth-defect. OGT catalyzes all nucleocytoplasmic O-GlcNAcylation, and we use unbiased MS-profiling to show that the short-term CDK12 inhibition induces hyper-O-GlcNAcylation of the spliceosome-machinery in PC and CRPC cells. Integration of DepMap- and a small scale-drug screen data reveled that depletion of CDK12 activity causes addiction to non-essential spliceosome components (CLK1/4 and SRPK1). CDK12-mutant tumors overexpress CLK1/4 and SRPK1. Finally, we show that the genomes of the CDK12-mutant tumors have lower DNA methylation, and that CDK12 inhibition induces the expression of the genes marked by DNA methylation.

**Conclusions:** Compromised CDK12 activity rewires DNA methylation, transcription and splicing, and this rewiring renders the affected cells addicted on the non-essential spliceosome components. We propose that inactivation of CDK12 is a biomarker for sensitivity against inhibitors of the non-essential spliceosome components just entering the clinical trials.

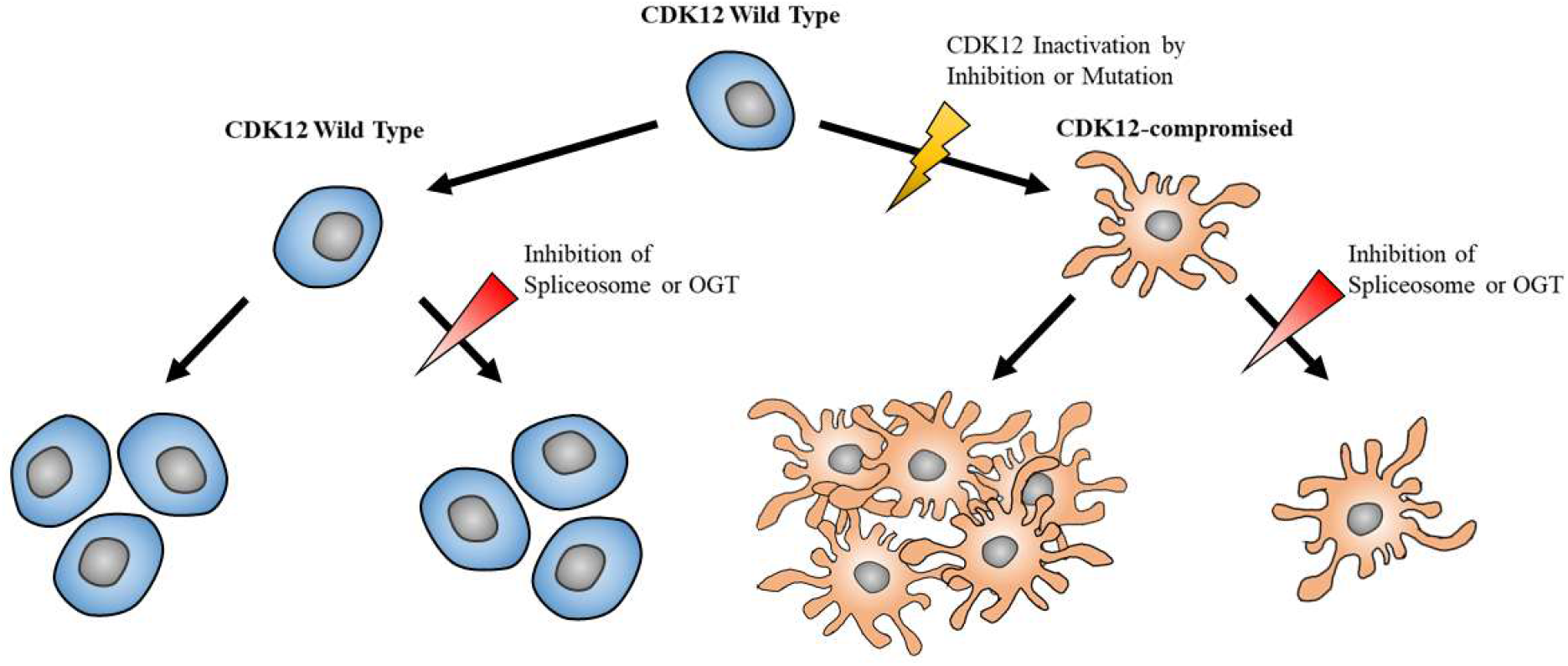

## Introduction

Prostate cancer (PC) is the most common cancer in men and after development of the castration-resistant PC (CRPC), there are no curative treatment options [1]. PC is typically a slow-progressing disease of the elderly men. Practically all tumors are positive for the nuclear hormone-activated transcription factor, androgen receptor (AR), and blocking the AR-activity halts disease progression [2]. CRPC occurs when the PC cells proliferate under the therapy-induced castrate-conditions. Many therapies are being trialed against this disease, and better understanding of the CRPC-specific features is likely to reveal points of vulnerability.

CRPC genomes accumulate mutations that support proliferation of the malignant cells, and it is these alterations that may be targeted as a CRPC-selective treatment-strategy. Specific mutations are enriched in the CRPC setting [3, 4]. Of a particular interest, inactivating mutations in the cyclin-dependent kinase 12 (CDK12) are 7-times more frequent in the CRPC when compared to primary tumors [5]. CDK12 mutations are rare in the primary PC, but when they occur, these tumors metastasize with almost 100% penetrance [6]. CDK12-mutant tumors define an aggressive sub-type of CRPC [7, 8], and by understanding the rewiring of these mutant-cells, we may discover actionable synthetic lethal interactions.

CDK12 phosphorylates RNA polymerase II (RNA Pol II) to suppress premature poly-adenylation and to support the transcription of the long genes [9]. RNA Pol II activity is also regulated by CDK7 and CDK9, to initiate the transcription and to release the polymerase for the productive transcription elongation, respectively. We discovered a PC cell-selective combinatorial lethal interaction between inhibitors of CDK9 and O-GlcNAc transferase (OGT) [10]. OGT catalyzes all nucleocytoplasmic O-GlcNAcylation of proteins, and we have shown that the enzyme is overexpressed in the aggressive PC [11, 12]. We hypothesized that better understanding of the crosstalk between OGT and CDK12 might enable discovery of a candidate treatment-strategy against the aggressive PC-sub-group, the CDK12-mutant tumors.

Here, we show that combined inhibition of OGT and CDK12 is selectively toxic to PC and CRPC cells, and that short-term CDK12 inhibition leads to an OGT-dependent remodeling of the splicing-machinery. Accordingly, CDK12 inhibition significantly alters the global splicing in PC cells. The effects on the global transcriptional program and splicing are mediated in part through DNA methylation, and CDK12-mutant tumors have decreased DNA methylation. We go on to show that co-targeting of CDK12 with five different spliceosome regulators is combinatorically lethal to CRPC cells, and, importantly, three of these spliceosomal proteins, CLK1, CLK4 and SRPK1, are non-essential. To conclude, our study proposes that compromised CDK12 activity render aggressive CRPC cells acutely reliant on the spliceosome, and we propose that CDK12-mutant tumors are susceptible to SRPK1 and CLK1/4 inhibitors, just entering the Phase 1 clinical trials.

## Results

### Combined inhibition of OGT and CDK12 is toxic to prostate cancer cells

CDK12 inactivating mutations are enriched in the aggressive CRPC [7, 8] but small molecule inhibitors targeting CDK12 halt the proliferation of prostate cancer cells [13]; clearly better understanding on how CRPC cells adapt to declining CDK12 activity is of high interest. We recently reported a combinatorial lethality screen that discovered a cancer cell-selective treatment combination using OGT and CDK9 inhibitors [10]. CDK9 is the transcription elongation kinase that releases RNA Pol II for the productive transcription elongation, while CDK12 maintains the phosphorylation in the long genes [9]. We hypothesized that we could enhance the anti-proliferative effects of CDK12 inhibition using OGT inhibitors. First, we treated prostate cancer cells with increasing doses of the CDK12 inhibitor THZ531 [14] in the presence and absence of the OGT inhibitor OSMI-4 [15]. OGT inhibition significantly and dose-dependently enhanced the anti-proliferative effects of the CDK12 inhibitor on two CRPC models (LN95 and 22RV1; **Fig. 1A**). Second, we selected a low dose of the CDK12 inhibitor and used live-cell imaging to follow the growth-kinetics of prostate cancer cells. Combining OGT and CDK12 inhibitors led to an almost complete growth-arrest of prostate cancer cells (LNCaP) and CRPC cells (LN95, C4-2 and 22RV1; **Fig. 1B**). Third, we measured cell death activation after CDK12 and OGT inhibitor treatments, which revealed a 20-fold increase in the PC and CRPC cells when compared to two cell lines representing normal prostate cells (PNT1 and RWPE-1; **Fig. 1C**).

**Figure 1.**
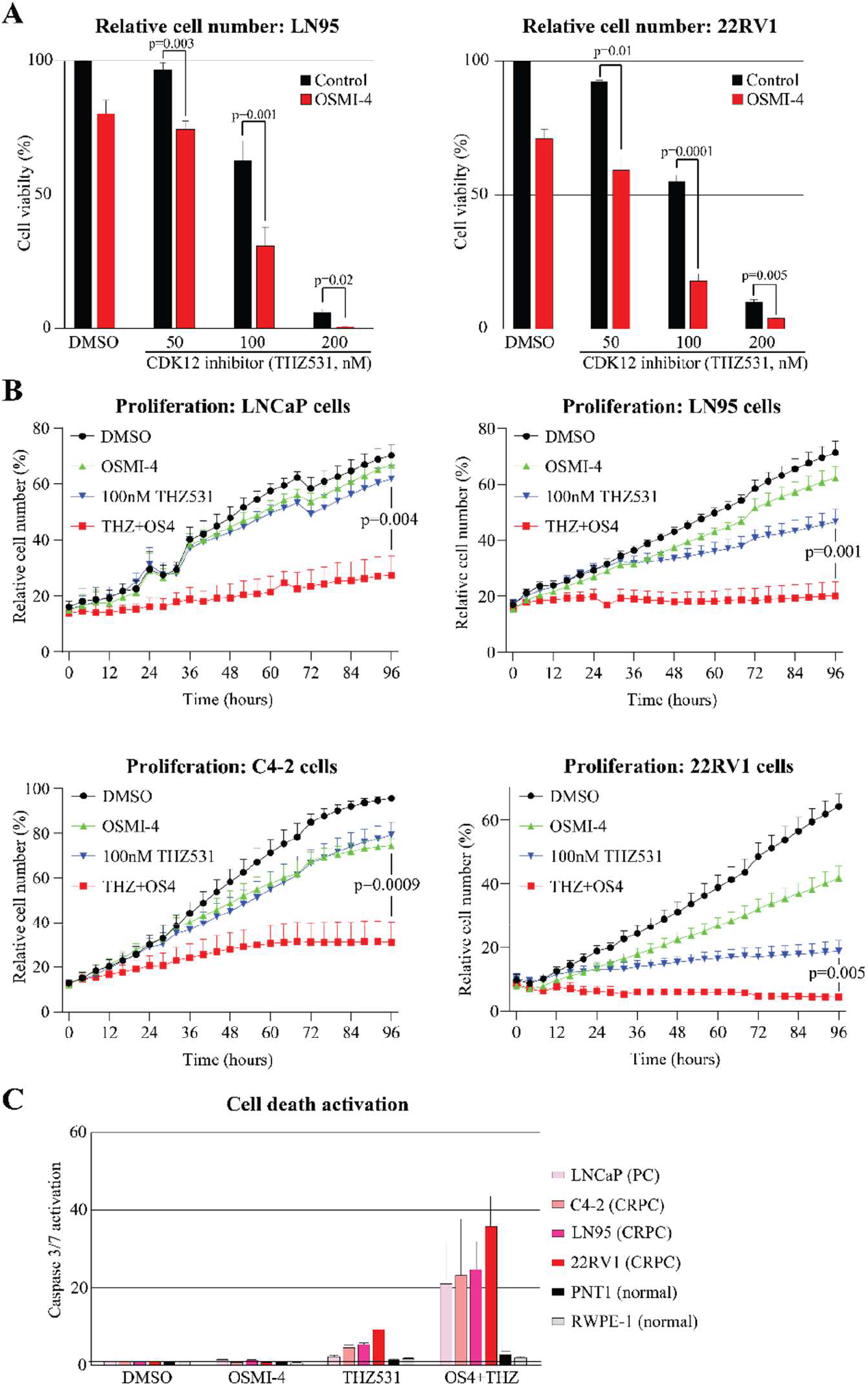
Combined inhibition of CDK12 and OGT is combinatorially lethal to prostate cancer cells. **A)** Cell viability of castration-resistant prostate cancer cells (CRPC, LN95 and 22RV1) was measured after four days of treatment using the CellTiterGlo-assay. Data shown is an average of four biological replicates with standard error of mean (SEM). Significance of the data was evaluated using the two-tailed paired Student’s t-test. **B)** Proliferation of PC cells (LNCaP) and CRPC cells (LN95, C4-2, 22RV1) was followed using live cell imaging every four hours. Data is presented as average of four biological replicates with SEM. Statistical analysis as in A. **C)** Cell death activation (caspase 3/7 cleavage) in PC, CRPC and normal prostate (PNT1 and RWPE-1) cells after four days of treatment (100nM THZ531). Data is presented as average of four biological replicates with SEM. DMSO sample was set to value of 1, and the other treatments are presented relative to that. OSMI-4 dose was 20μM in all experiments of figure 1.

Combined targeting of OGT and CDK12 robustly suppresses the proliferation of prostate cancer cells but we did not yet know why this is. It is well-established that OGT activity is remodeled in response to stress [11, 16]. Therefore, we hypothesized that by identifying the proteins OGT modifies when CDK12 activity is depleted, we may be able to design a rational drug-combination that selectively targets the cells with compromised CDK12 signaling.

### CDK12 inhibition induces hyper-O-GlcNAcylation of the spliceosome machinery

We used mass spectrometry to identify the proteins that get hyper-O-GlcNAcylated after short-term treatment with the CDK12 inhibitor. We performed anti-O-GlcNAc immunoprecipitation experiments followed by mass spectrometry analysis in biological triplicates in PC and CRPC cells (LNCaP and LN95). LN95 cell line was established through an extended androgen-deprivation of the androgen-dependent LNCaP cell line. Combined inhibition of OGT and CDK12 is toxic to both PC and CRPC cells (**Fig. 1**), so we hypothesized that by using an isogenic system that models the development of the CRPC, we will identify the critical factors that are O-GlcNAcylated when CDK12 is inhibited. As expected due to the short treatment time, most of the O-GlcNAcome remained intact, but there were a number of proteins that were increasingly modified by OGT when CDK12 was inhibited (**Fig. 2A**). We selected the proteins whose O-GlcNAcylation increased at least 20% in all of the biological replicates and performed pathway enrichment analysis, which revealed the spliceosome as the most enriched category (**Fig. 2B**). Based on these data, CDK12 inhibition induces an OGT-dependent remodeling of the spliceosome activity.

**Figure 2.**
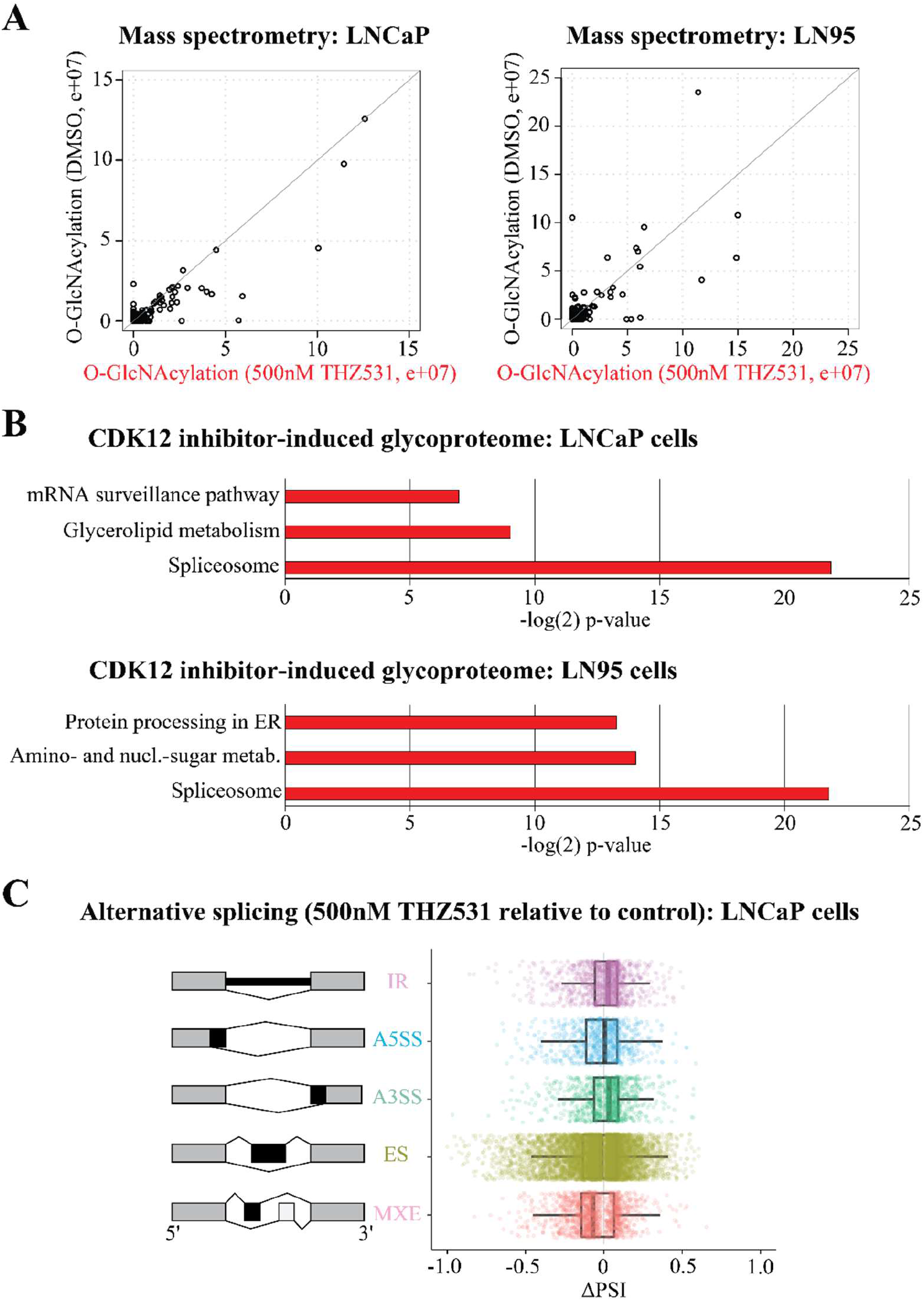
Inhibition of CDK12 induces OGT-dependent remodeling of the spliceosome. **A)** Scatter plot of proteins O-GlcNAcylated in response to CDK12 inhibition. LNCaP and LN95 cells were treated for 4 hours with 500nM THZ531, after which O-GlcNAcylated proteins were enriched using immunoprecipitation and analyzed by mass spectrometry. Data shown is an average of three biological replicates, which were first normalized to control-immunoprecipitation. **B)** KEGG-pathway enrichment analysis of proteins whose O-GlcNAcylation increased by 20% after THZ531 treatment in all three biological replicates, when compared to control cells. **C)** CDK12 inhibitor-induced alternative splicing (AS). Left are the major alternative splicing events: IR: intron retention; A5SS: alternative 5′ splice sites; A3SS: alternative 3′ splice sites; ES: exon skipping; MXE: mutually exclusive exon. Right: AS-events in LNCaP cells treated with 500nM THZ531 for 6 hours relative to DMSO-treated control.

CDK12 has a well-established function as the transcription elongation kinase whose activity is important to suppress premature poly-adenylation [9], but it is currently not known if CDK12 inhibition affects splicing in prostate cancer cells. We therefore employed the rMATS algorithm [17] to assess if short-term (6 hours) inhibition of CDK12 activity affects global splicing by measuring alternative 3′ and 5′ splice sites (A3’SS and A5’SS, respectively), mutually exclusive exons (MXE), detained introns (DI) and skipped exons (SE) (**Fig. 2C**). We detected 10 000 significant alternative splicing events by comparing CDK12 inhibitor treatment to control treatment (**Fig. 2C**). Particularly exon definition (MXE and SE) was affected. Of a particular interest, processing of the androgen receptor mRNA was faulty due to SE (p=4.17e-08 as calculated in the rMATS-analysis). When we visualized RNA-seq data for the AR locus, we noted a robust increase in the reads of the cassette exon 3 for the AR mRNA in all of the CDK12 inhibitor treated samples (**Suppl. Fig. 1A**). Inclusion of this exon introduces a premature stop codon, and leads to generation of a ligand-independent AR, AR-v7 [18]. AR-v7 expression emerges in response to anti-androgen therapy and is associated with development of the CRPC disease [19]. It is possible that the compromised CDK12 activity, as observed in the aggressive CRPC, promotes generation of AR-v7 splice variant. One observation supporting this hypothesis is that CDK12 mutant tumors express higher levels or AR-v7 (**Suppl. Fig. 1B**).

To summarize our findings so far, CDK12 inhibition increases O-GlcNAcylation of the spliceosome machinery in PC and CRPC cells, and additionally affects global splicing. We hypothesized that the CDK12 inhibitor-induced transcriptional stress renders CRPC cells addicted on the specific spliceosomal proteins.

### Inhibition of CDK12 induces dependency on the non-essential components of the spliceosome

We moved on to target the spliceosome in combination with CDK12 inhibitor to identify the combinatorial lethal partners. Spliceosome is a complex machinery, and only the major druggable steps are described here [20]: First, U1 snRNP binds to the 5’ end of the pre-mRNA and other non-snRNP associated factors, to form the commitment complex (**Fig. 3A**). Second, U2 snRNP binds to the branch region to form complex with SF3B. SF3B is part of the core splicing factors and it is required to form the stable complex between U2 snRNP and the pre-mRNA; both of which are required for processing of most of the mRNAs [20]. So-called SR-proteins are the major components affecting both constitutive and the alternative pre-mRNA splicing. In addition, SR proteins also regulate mRNA export, genome stability, nonsense-mediated decay and translation. The name of the SR-proteins comes from the long repeats of serine (S) and arginine (R) amino acids: serines are extensively phosphorylated by SRPKs and CLKs, while arginines are methylated by the PRMT-proteins. These post-translational modifications have gene specific effects, and inhibition of CLK1, CLK4 or PRMT5 activities selectively affects splicing of the different sub-sets of mRNAs [21, 22].

**Figure 3.**
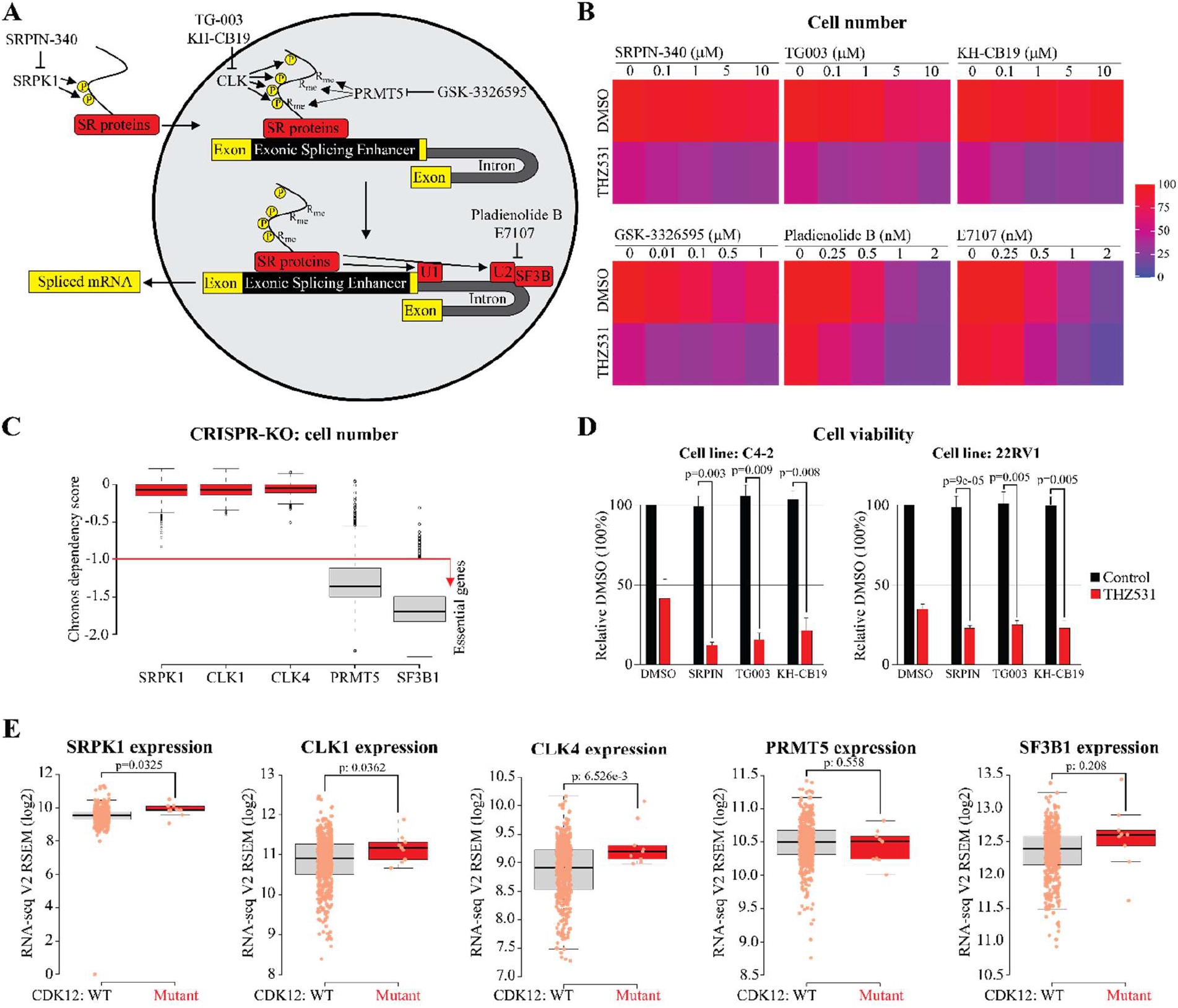
Compromised CDK12 activity causes dependency on the non-essential spliceosome components. **A)** Targeting spliceosome with small molecule inhibitors. **B)** Heat map of 22RV1 cell number after four-day treatment with spliceosome inhibitors either in the presence or absence of 50nM THZ531. The relative cell number was measured using crystal violet-assay. Data represents average of 2-3 biological replicates normalized to DMSO. **C)** CRISPR-mediated depletion of SRPK1, CLK1 and CLK4 has minimal effect on cell number, while depletion of PRMT5 and SF3B1 has a highly significant effect. The plot was generated using the data available through the DepMap CRISPR-database (DepMap 21Q4 Public+Score, Chronos) [23]. The score of 0 indicates that the gene is non-essential, and score of -1 (highlighted in red here) is the median of all pan-essential genes. **D)** Viability (CellTiterGlo) of CRPC cells after 3 days treatment. Data shown is an average of 3-4 biological replicates with SEM. Significance of the data was evaluated using the two-tailed paired Student’s t-test. **E)** SRPK1, CLK1 and CLK4 are significantly overexpressed in the CDK12-mutant tumors when compared to CDK12-wild type tumors. The plot was generated using the TCGA dataset [24] accessed through the cBioPortal [41, 42]. Significance was reported in cBioPortal and represents Student’s t-test.

CDK12 inhibition causes increased dependency on the spliceosome. Specific compounds targeting SRPK1, CLK1 / CLK4, PRMT5 and SF3B1 have been described, and here we used these compounds to establish potential combinatorial lethal interactions with CDK12 inhibition (**Fig. 3A**). All of the spliceosome inhibitors sensitized CRPC cells to CDK12 inhibitor (**Fig. 3B**). Remarkably, inhibitors of SRPK1 and CLK1 / CLK4 did not have any effect as a single agent up to 10μM dose, but robustly sensitized the CRPC cells to the CDK12 inhibitor THZ531. On the other hand, PRMT5 and SF3B1 inhibitors were toxic also as a single agent treatment.

The lack of a single-agent toxicity for the SRPK1 and CLK1 / CLK4 inhibitors implies that these factors are non-essential in most cell types. We used the DepMap comprehensive CRISPR-KO database to assess this [23]. As expected based on the single agent toxicity, SF3B1 and PRMT5 are essential genes in most cell types (**Fig. 3C**). Interestingly, however, depletion of SRPK1, CLK1 or CLK4 minimally affected the proliferation of the cells: these three factors are redundant in most cell types and represent ideal targets for synthetic lethal interactions.

We focused on SRPK1, CLK1 and CLK4 as the candidate combinatorial lethal targets with the compromised CDK12 activity. We confirmed that inhibition of these factors sensitizes another CRPC model (C4-2) to CDK12 inhibitor THZ531 using viability and cell proliferation assays (**Fig. 3D** and **Suppl. Fig. 2**). To conclude, depletion of CDK12 activity sensitizes CRPC cells to SRPK1 and CLK1/4 inhibitors.

We hypothesized that the expression of the splicing-regulatory factors is altered in the CDK12-mutant tumors. To test this, we identified CDK12-mutant tumors in primary prostate cancer using the TCGA dataset [24], and discovered that the expression of SRPK1, CLK1 and CLK4 was significantly increased in the CDK12 mutant tumors (**Fig. 3E**). In contrast, the expression of PRMT5 and SF3B1 was not significantly different between the wild type and the CDK12-mutant tumors.

To summarize our findings so far, decline in the CDK12 activity sensitizes CRPC cells to inhibitors of SRPK1, CLK1 and CLK4, and CDK12-mutant tumors overexpress these three non-essential spliceosome-regulators. However, we did not yet know how the global transcription is rewired to sustain the aggressive phenotype when the activity of a major transcriptional kinase, CDK12, is compromised.

### Compromised CDK12 activity induces the expression of genes marked by DNA methylation

In order to understand how the decrease in CDK12 activity affects global transcription, we generated gene expression profiles after CDK12 inhibitor treatment. As previously reported, CDK12 inhibition robustly decreases the expression of mRNAs related to DNA repair [25], and we used these as positive controls here (highlighted in red, **Fig. 4A**). Unexpectedly because CDK12 is a major transcriptional kinase, we noted that 38% of the significantly affected mRNAs were increased in their relative abundance in response to CDK12 inhibition.

**Figure 4.**
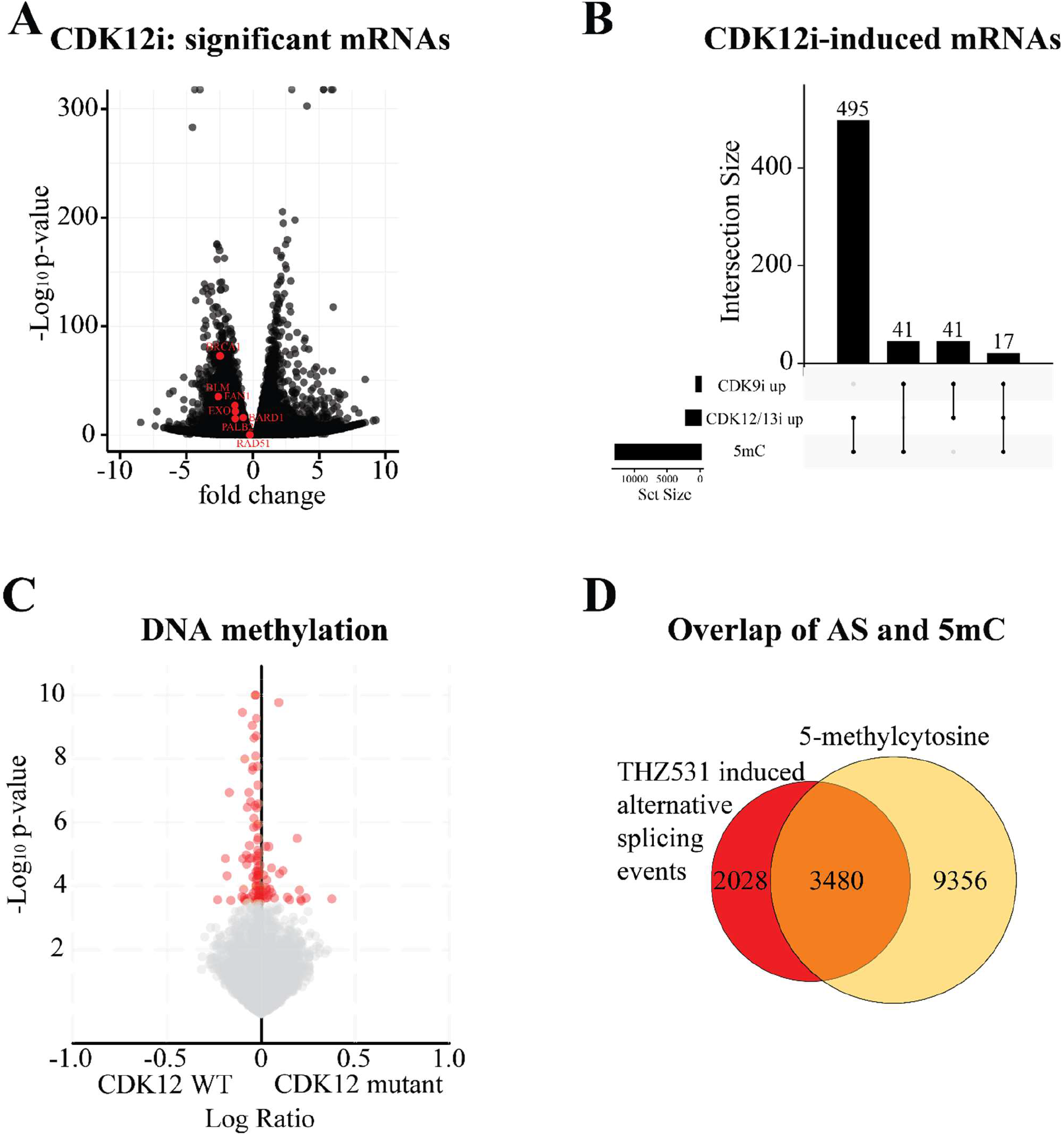
Compromised CDK12 activity induces the expression of genes marked by DNA methylation. **A)** CDK12 inhibitor-induced effects on gene expression. Volcano plot shows mRNAs whose expression change significantly (p<0.05) in LNCaP cells treated with 500nM THZ531 for 6 hours relative to DMSO-treated control. Genes highlighted in red are those previously reported [25] to be downregulated in response to CDK12 inhibitor treatment. **B)** CDK12 inhibition induces mRNAs that are marked by 5-methylcytosine (5mC). Upset-plot presenting the overlap between the genes marked by 5mC and mRNAs induced by CDK12 inhibition (p<0.05), and mRNAs induced by CDK9 inhibition (p<0.05). **C)** CDK12-mutant tumors have less DNA methylation than the CDK12-wildtype tumors. The plot was generated using the tools available in cBioPortal [41, 42] and the data is from the TCGA dataset [24]. **D)** A high number (63%) of CDK12 inhibitor-induced alternative splicing events are found on the genes that are marked by 5mC. Alternative splicing events are from **Fig. 2C** and 5mC data from **Fig. 4B**.

We assessed if the typical epigenetic marks are enriched on the genes induced by the CDK12 inhibitor. None of the canonical histone marks (h3K4me1, h3K4me3 or h3K27me3) were enriched in the promoters of these genes (**Suppl. Fig. 3A**). In contrast, we noted that a high number of the CDK12 inhibitor-induced genes were marked by DNA methylation (25%, **Fig. 4B**). DNA methylation is typically associated with the lack of gene expression, but can be removed through active demethylation [26]. We therefore assessed if the enzyme removing DNA methylation or the de-methylation intermediate, 5-hydroxymethylcytosine, are found on the CDK12 inhibitor-induced mRNAs. However, we did not observe a robust overlap with either (**Suppl. Fig. 3B**).

DNA methylation is an abundant epigenetic mark, and we therefore assessed if the mRNAs increased in response to CDK9 inhibitor treatment are also marked by DNA methylation, but they are not (**Fig. 4B**). Finally, we asked if CDK12-inactivation through mutation alters DNA methylation in the patient samples. Indeed, CDK12-mutant tumors have less DNA methylation (**Fig. 4C**). Based on these data, we propose that the compromised CDK12 activity alters the transcriptional program in part by affecting the genes marked by DNA methylation.

DNA methylation is known to regulate splicing [27], and we next focused on the crosstalk between CDK12 inhibition, splicing and DNA methylation. We noted that 63% of the CDK12 inhibitor-induced alternative splicing-events coincide with DNA methylation (**Fig. 4D**). Two different primary mechanisms convey information from DNA methylation to regulate alternative splicing [27]. In the first, RNA polymerase II velocity is regulated via CCCTC-binding factor (CTCF) and methyl-CpG binding protein 2 (MeCP2). In the second, heterochromatin protein 1 (HP1α, HP1β or HP1γ) binds to methylated DNA and recruits the splicing machinery. We therefore asked if the CDK12-mutant tumors overexpress any of these factors. HP1γ was the only gene significantly overexpressed in the CDK12-mutant tumors (**Suppl. Fig. 4A, B**). Next, we turned to our mass spectrometry data to assess if CDK12 inhibition affects O-GlcNAcylation of HP1γ, and noted that it was increasingly O-GlcNAcylated in both LNCaP and LN95 cell lines in response to CDK12 inhibition (**Suppl. Fig. 4C**). We propose that OGT regulates HP1γ to coordinate with the splicing machinery when CDK12 activity is compromised, but functional elucidation of this requires a separate, comprehensive study.

To conclude, here we show that the compromised CDK12 activity remodels global splicing in an OGT-dependent manner. Our study discovered a process that links CDK12 and DNA methylation to splicing. Finally, we propose that inactivation of CDK12 is a biomarker for sensitivity against SRPK1 and CLK1/4 inhibitors, which are currently evaluated in clinical trials.

## Discussion and conclusions

Here, we discovered a combinatorial lethal interaction between compromised CDK12 activity and the non-essential components of the spliceosome. This project was initiated based on two notions: defects in CDK12 define an aggressive subtype of prostate cancer [3-6], and combined inhibition of OGT and CDK9, a closely functionally related kinase to CDK12, induces cytotoxic effects [10]. Development of the highly specific inhibitors against both OGT and the transcriptional kinases enables direct dissection of the crosstalk between phosphorylation and O-GlcNAcylation.

We show that the decrease in the CDK12 activity leads to an OGT-dependent adaptive remodeling of the spliceosome, and spliceosome is a point of vulnerability when CDK12 is compromised. First, we confirmed that the combined inhibition of OGT and CDK12 causes combinatorial lethal interaction (**Fig. 1**). Second, we used an unbiased O-GlcNAcome profiling to discover that CDK12 inhibition directs OGT to O-GlcNAcylate components of the spliceosome (**Fig. 2B**). OGT has been previously linked to splicing [28-30], and here we show that the high OGT activity becomes critical when CDK12 activity is compromised. Third, we revealed that CDK12 inhibition induces alternative splicing (**Fig. 2C**). This is an interesting notion, as it has been established that aggressive prostate cancer tumors are characterized by defects in the spliceosome machinery [31]. Based on our data, we hypothesize that the altered CDK12 activity should be mutually exclusive with alterations in the spliceosome genes, but this remains to be determined. Fourth, we showed that inhibition of CDK12 activity renders CRPC cells dependent on the non-essential components of the spliceosome (**Fig. 3B and 3C**). Our work proposes CDK12 inactivation as a biomarker for the SRPK1 and CLK inhibitors just entering the clinical trials. Fifth, we showed that the compromised CDK12 activity leads to increased expression of a high number of mRNAs that are marked by DNA methylation (**Fig. 4B**). It was unexpected that inhibition of the major transcriptional kinase increases the expression of a high number of genes. This effect appears to depend on the cell type, and in certain models, CDK12 inhibition decreases the expression of most of the mRNAs, while in others, a high number of mRNAs is increased [13, 25, 32]. The apparent increase in certain mRNAs is likely explained in part due to the premature polyadenylation and increased transcription initiation: It is known that CDK12 inhibition causes premature poly-adenylation [9]. In support of the latter, we noted that targeting CDK12 increases transcription initiation of the androgen receptor-gene, causes inclusion of an alternative exon and lower number of reads towards the end of the gene (**Suppl. Fig. 1**). Neither the existing literature, nor the data presented here, can thoroughly explain how CDK12 affects global DNA methylation and splicing to allow the maintenance of the pro-proliferative transcriptional program. Nevertheless, compromised CDK12 activity renders the CRPC cells dependent on the non-essential spliceosome-regulators.

Integration of the genome-wide CRISPR knockout data and compound-screening data represents an appealing strategy to identify low-risk synthetic lethal partners. Normal and cancer cells utilize the same machinery to carry out their ‘functions’, which in the case of cancer cells is to produce more cells. In practical terms, this means that the majority of the cancer therapies have severe on-target, undesirable side effects. Cancer cell-specific genetic alterations are a characteristic that segregates the malignant cells from the normal. This has led to selective and truly life-changing therapies, one of the most famous examples being Imatinib, the compound that blocks the proliferation of BCR-ABL fusion-dependent cells [33]. Unfortunately, truly actionable mutations are rare, and scientific community often relies on the existing therapies and uses the mutations discovered through high-throughput sequencing as biomarkers of the hypothesized sensitivity.

Compromised CDK12 activity lowers the DNA repair efficiency, and CDK12 mutations have therefore been proposed as a biomarker for PARP inhibitors and immunotherapy [6-8, 34]. Decreased genome stability of the CDK12-mutant tumors is in the epicenter of both of these treatment strategies, and the current model proposes that the altered CDK12 activity causes DNA damage due to decreased expression of the DNA repair factors, and we also noted decreased expression of a number of the DNA repair factors when CDK12 is inhibited (**Fig. 4A**). However, CDK12 inhibitors induce DNA damage rapidly, while the loss of DNA repair factor expression first occurs at the transcriptional level and, eventually, at the protein level. It appears that the CDK12 inhibitor-induced DNA damage is explained through a more complex mechanism that may involve collisions with the replication machinery and / or loss of the coordination between transcription, DNA methylation and splicing, and here we provide evidence for the latter. Decrease in CDK12 activity leads to a lowered ability to repair DNA, and in combination with the sustained DNA damage, causes large-scale genomic alterations, as observed in the CDK12-mutant tumors [5, 35]. Understanding functionally, why the compromised CDK12 activity causes genome instability is an important goal for the future research.

To conclude, here we report a novel combinatorial lethal interaction between the compromised CDK12 activity and the non-essential components of the spliceosome. Our study shows how OGT activity is rapidly remodeled in response to decrease in the CDK12 activity. Applying similar approach to other treatments is a viable strategy to identify functionally relevant targets for other post-translational modifications as well. In the future, it is important to dissect the signals / factors that direct OGT activity in response to stimuli.

## Methods

### Cell culture, compounds and preparation of cell lysates

LNCaP, 22RV1, C4-2, and RWPE-1 cell lines were obtained from the American Tissue Culture Collection (ATCC) and PNT1 cell line was acquired from Sigma. LN95 cell line was kindly provided by Professor Stephen Plymate (University of Washington). LNCaP, 22RV1, C4-2 and PNT1 cells were maintained in RPMI medium supplemented with 10% fetal bovine serum (FBS). RWPE-1 cells were cultured in keratinocyte serum-free media, while LN95 cells were maintained in phenol red-free RPMI supplemented with 10% charcoal-stripped FBS. All cell lines were allowed to adhere for at least one day prior to the treatment with inhibitors. THZ531, OSMI-4, TG003 and GSK3326595 were obtained from MedChemExpress. KH-CB19, Pladienolide B and SRPIN340 were purchased from Tocris.

### Proliferation assays

Proliferation rate of cells was measured using Incucyte live cell imaging system following manufacturer’s (Sartorius) instructions. For viability assays, we used CellTiter-Glo® (only in **Fig. 1A**) and CellTiter-Glo® 2.0 assays (Promega). To detect the cell death activation, we used IncuCyte Caspase-3/7 Green Reagent for apoptosis (Sartorius). Crystal violet staining assay was used to assess the relative number of cells as previously described [36], with minor changes: Cells growing on a 96-well plate were fixed first with ice-cold 70% methanol for 2 minutes and then with 100% methanol for 10 minutes. Cells were allowed to dry after which they were stained with 0.05% crystal violet solution for 10 minutes and washed twice with deionized water. After the cells dried out, 50μL of 10% acetic acid was added to each well, the cells were de-stained for 15 minutes on a plate shaker and the absorbance was measured.

### Mass spectrometry

For O-GlcNAc immunoprecipitation, we combined two antibodies: RL2 (Abcam: ab2739) and CTD110.6 (Cell Signaling Technologies: 9875), and used Pierce Direct IP Kit (ThermoFisher Scientific) according to manufacturer’s protocol to prepare the biological triplicate samples for mass spectrometry (MS). MS was performed by the Weill Cornell Medicine (WCM) Meyer Cancer Center Proteomics & Metabolomics Core Facility.

### Bioinformatics

The RNA-seq data of LNCaP cells treated with 500nM THZ531 was downloaded from BioProject (Accession: PRJCA004744) [37]. First, the raw fastq files were aligned to human genome HG38 using STAR aligner [38]. Second, the aligned SAM files were converted into their binary formats (BAM) using Samtools [39]. Finally, we used DESeq2 [40] to call the differentially expressed genes from the BAM files. Threshold for p-Value of 0.05 and log fold change of either 1 or -1 was applied to define a gene as significantly up-or down-regulated. To identify the alternatively spliced sites we used rMATS v4.1.1 [17]. rMATs was run with genome index generated from STAR aligner. The “--variable-read-length” parameter was used to allow the processing of reads with read length different than the average read length. The essentiality and expression level of spliceosome components was assessed using data available through the DepMap CRISPR-database (DepMap 21Q4 Public+Score, Chronos) [23] and cBioPortal [41, 42] (TCGA dataset [24]), respectively. Boxplots were made using R version 4.1.1 in R studio 2021.09.0. The volcano and boxplots for the transcriptomics data were made in R Studio. The ChIP-seq for H3Kme1, H3Kme4 in LNCaP cells was downloaded from GSE14092 [43] and data for H3K27me3 was obtained from GSE107780 [44]. We used the genome wide occupancy data for TET2, 5-methylcytosine (5mC) and 5-hydroxymethylcytosine (5hmC) deposited with GSE66039 accession ID [44]. CDK9 inhibitor induced mRNAs in LNCaP cells treated with CDK9 inhibitor were obtained from [10]. Upset plot was generated using UpSetR package in R [45].

## Competing Interests

The authors have declared that no competing interest exists.

## Funding

HMI is grateful for the funding from the Academy of Finland (Decision nr. 331324 and nr. 335902) and the Jenny and Antti Wihuri Foundation.

## Authors’ contributions

SP performed the combinatorial compound testing of CDK12 and spliceosome inhibitors, evaluated spliceosome mRNAs in patient samples, evaluated DepMap-data and wrote the methods and figure legends. AG performed all of the bioinformatics analysis (alternative splicing, correlation of genesets with epigenetic features and made figures from the mass spectrometry data). DYD and CT provided reagents and feedback on the project. ML provided resources and critical feedback on the development of the project. HMI designed and supervised the study, and wrote the manuscript. All authors have reviewed the manuscript.

## Acknowledgements

The authors wish to acknowledge CSC – IT Center for Science, Finland, for the computational resources. Mass spectrometry was performed by the WCM Proteomics and Metabolomics Core Facility. Preliminary data for this project was generated in Professor Suzanne Walker’s laboratory (Harvard Medical School, supported by National Institutes of Health grant R01 GM094263). E7107 was a gift from H3Biosciences/Esai.

## Supplementary figures

**Supplementary figure 1.**
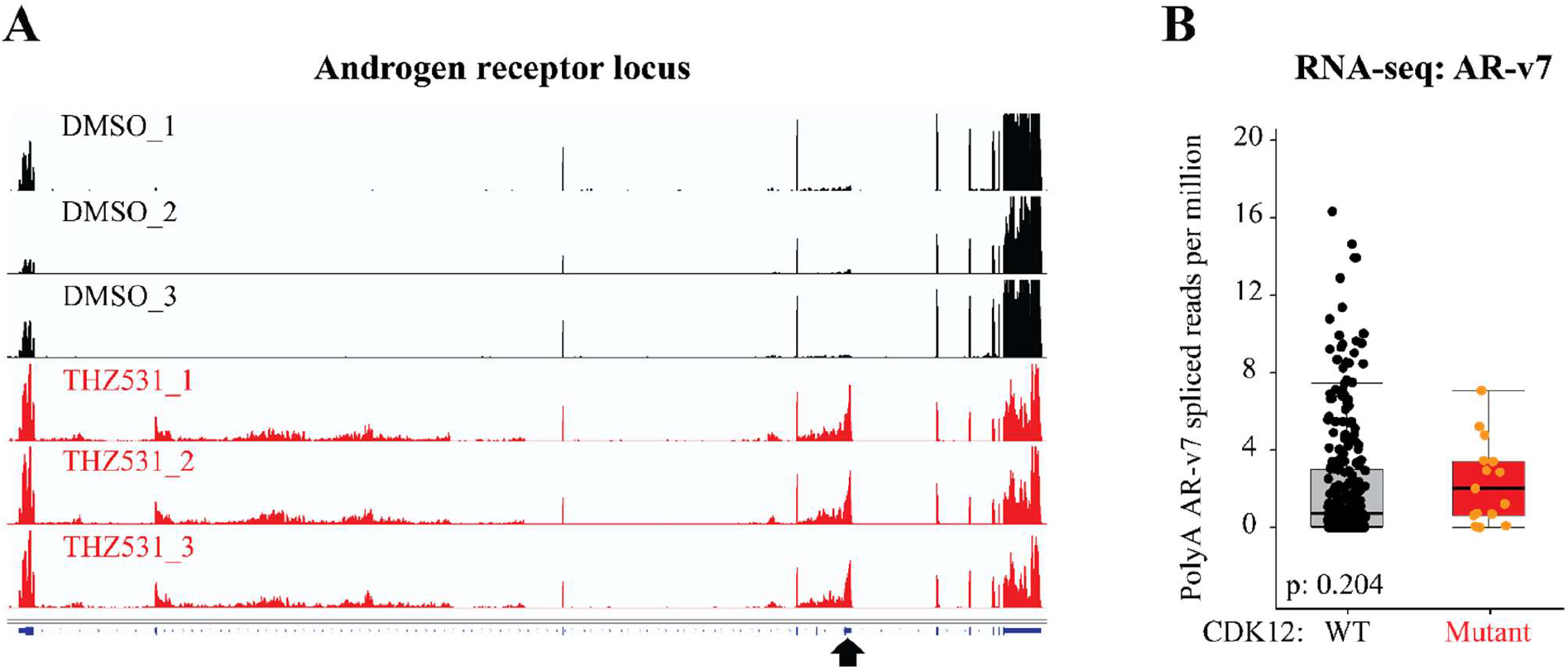
Compromised CDK12 activity affects processing of the androgen receptor (AR) mRNA. **A)** LNCaP cells treated with 500nM THZ531 for 6 hours show increased reads on an alternative exon highlighted with an arrow (data analyzed from previous publication: PRJCA004744 [37]). Inclusion of this exon leads to the generation of a ligand-independent form of AR, AR-v7. Y-axis was set to same value for all treatments. **B)** CDK12-mutant tumors express higher levels of AR-v7 when compared to CDK12-wild type tumors. The plot was generated using the SU2C/PCF Dream Team-dataset [4] accessed through the cBioPortal [41, 42]. Statistical analysis is the value reported in cBioPortal (p-value derived from Wilcoxon test).

**Supplementary figure 2.**
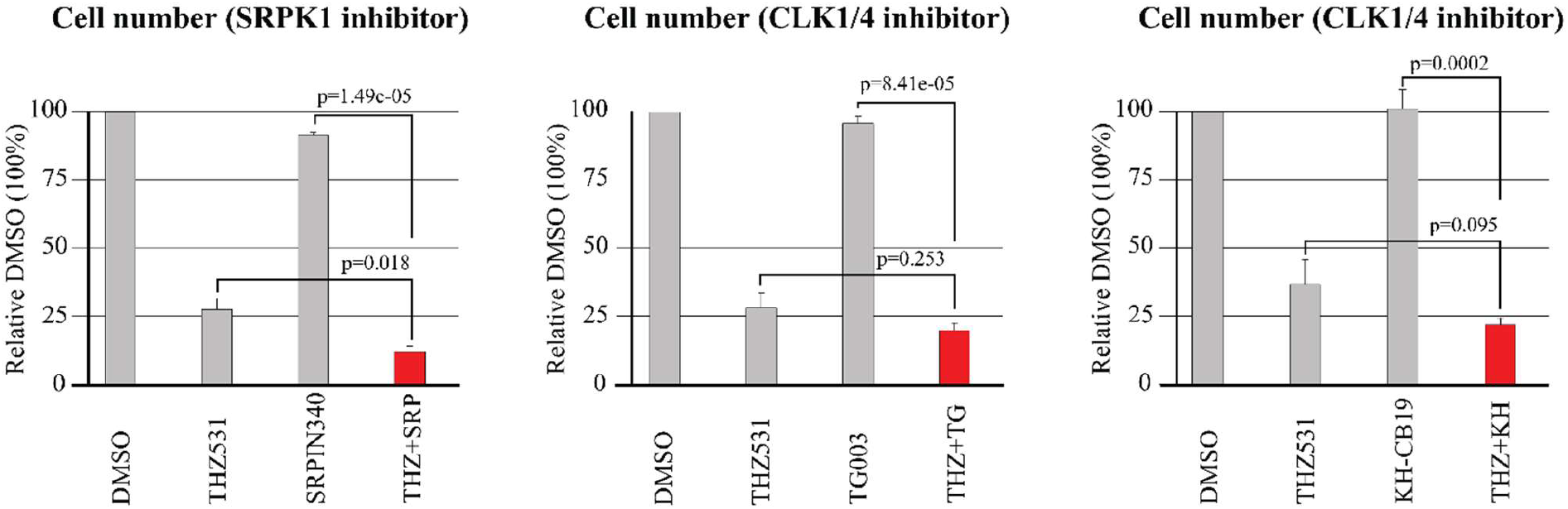
Combined inhibition of CDK12 and spliceosome reduces prostate cancer cell number. Bar plots showing the C4-2 cell number based on crystal violet-assay, and after four-day treatment with 100nM THZ531, 5μM spliceosome inhibitor (SRPIN340, TG003, or KH-CB19), or the combination of both. Data is presented as average of five biological replicates with SEM. Statistical analysis was performed using the two-tailed paired Student’s t-test.

**Supplementary figure 3.**
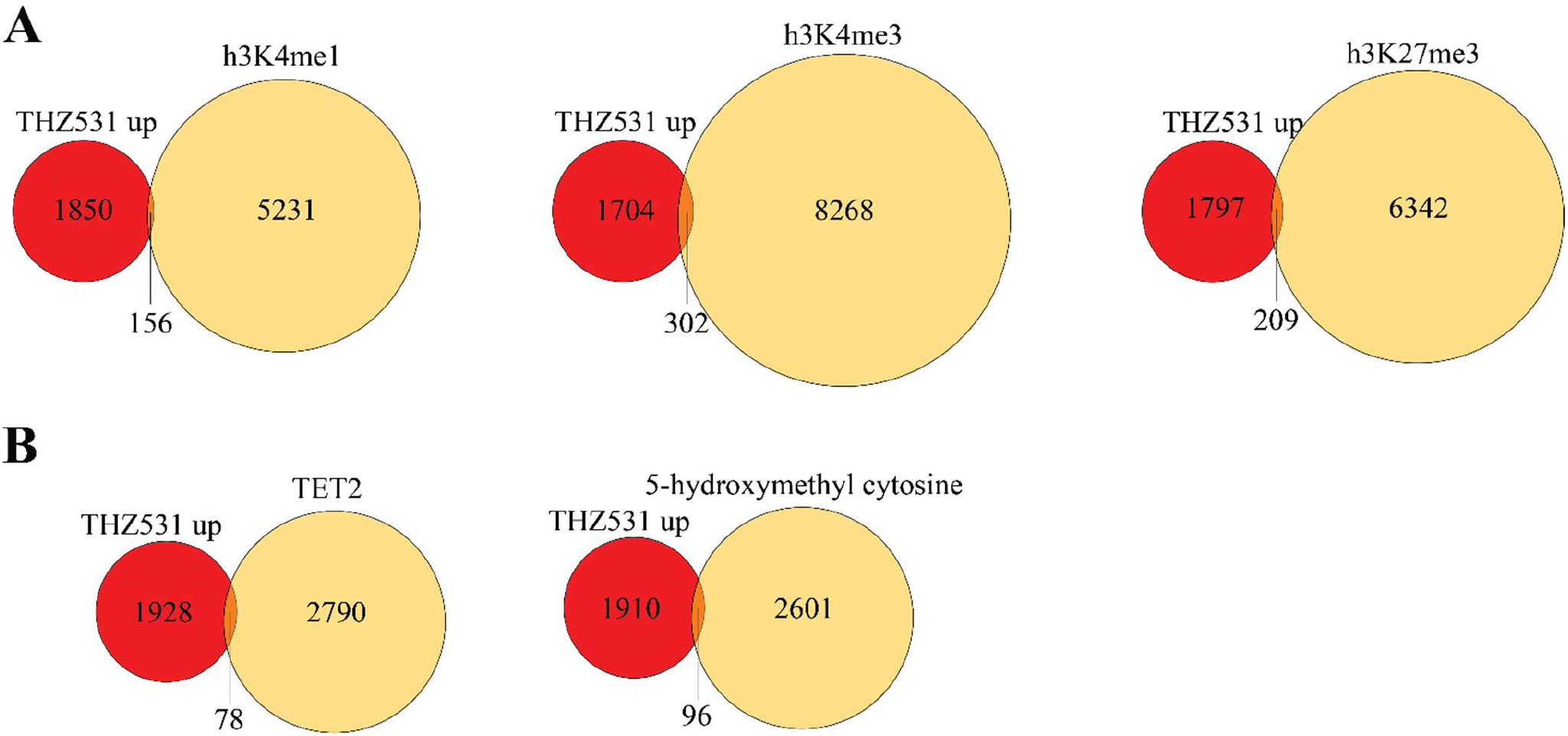
Overlap between CDK12 inhibitor-induced mRNAs and epigenetic marks. Datasets used are described in the methods section (THZ531 up represents mRNAs whose expression was significantly increased, p<0.05). **A)** Overlap between CDK12 inhibitor-induced mRNAs and the histone post-translational modifications. **B)** Overlap between CDK12 inhibitor-induced mRNAs and TET2 (left) or 5-hydroxymethyl cytosine (right).

**Supplementary figure 4.**
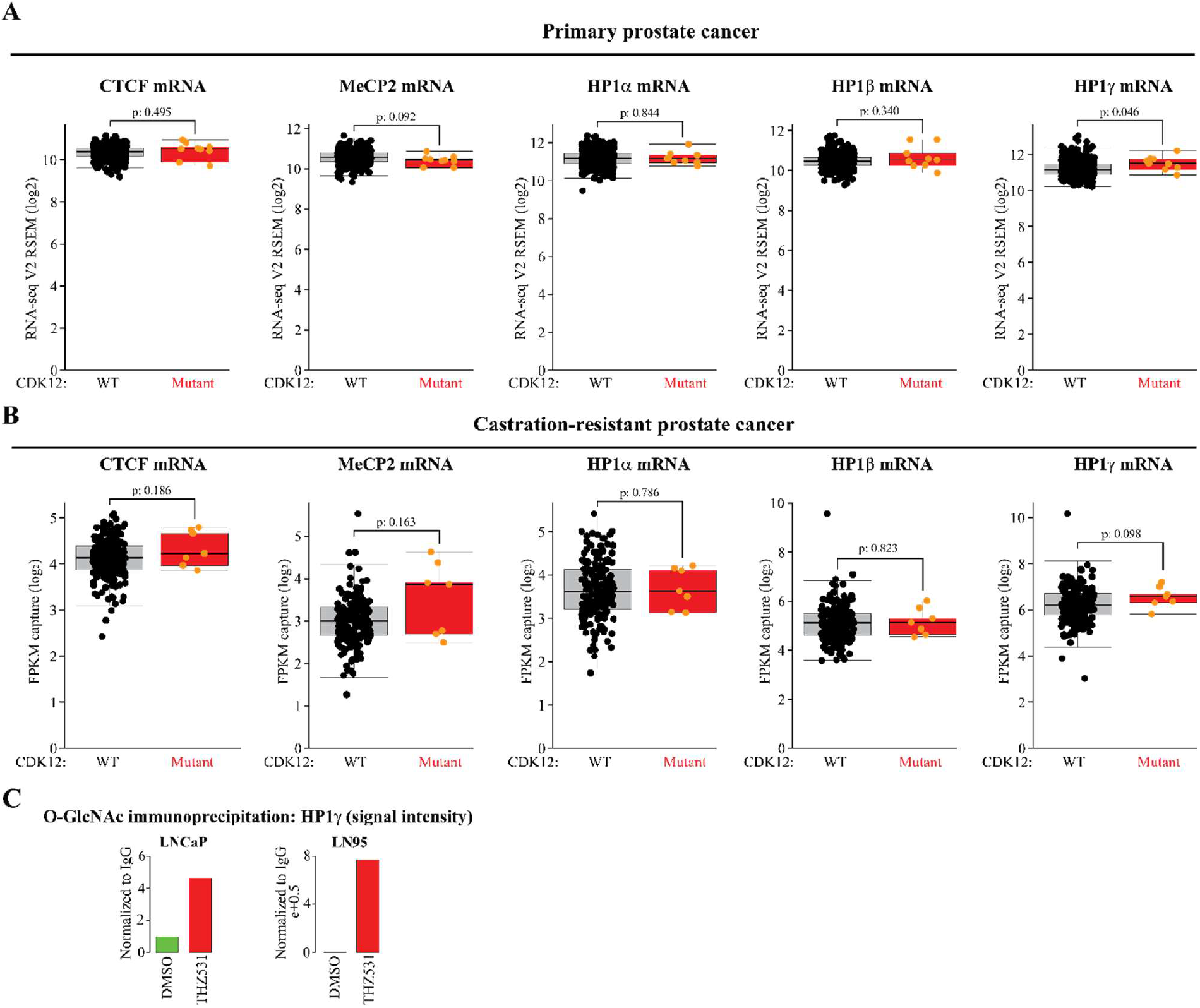
CDK12-mutant prostate cancers overexpress HP1γ. **A)** HP1γ is significantly overexpressed in the CDK12-mutant primary tumors when compared to CDK12-wild type tumors. The plot was generated using the TCGA dataset [24] accessed through the cBioPortal [41, 42]. Significance was reported in cBioPortal and represents t-test. **B)** HP1γ is overexpressed in the CDK12-mutant castration-resistant prostate cancer tumors when compared to the CDK12-wild type tumors. The plot was generated using the SU2C/PCF Dream Team-dataset [4] accessed through the cBioPortal [41, 42]. Significance was reported in cBioPortal and represents t-test. **C)** CBX3 is increasingly O-GlcNAcylated in response to 4 hours treatment with 500nM THZ531. The data shown is an average of three biological replicate immunoprecipitation experiments analyzed using mass spectrometry. Signal intensity was first normalized to negative immunoprecipitation and the data is presented as relative fold-change to DMSO, which was set to value of 1.

## Notes

### Competing Interest Statement

The authors have declared no competing interest.

